# Multimodal disease transmission as a limiting factor for the spatial extent of a host plant

**DOI:** 10.1101/2022.05.11.491512

**Authors:** Lawrence H. Uricchio, Emily L. Bruns, Michael Hood, Mike Boots, Janis Antonovics

## Abstract

Theoretical models suggest that infectious diseases could play a substantial role in determining species’ ranges, but few studies have collected the empirical data required to test this hypothesis. Pathogens that sterilize their hosts or spread through frequency-dependent transmission could have especially strong effects on the limits of species’ distributions because sterilized hosts can serve as long-lived disease reservoirs and frequency-dependent transmission mechanisms are effective even at very low population densities. We collected spatial disease prevalence data and population abundance data for alpine carnations infected by the sterilizing pathogen *M. violaceum*, a disease that is spread through both frequency-dependent (vector-borne) and density-dependent (aerial spore transmission) mechanisms. Our 13-year study reveals rapid declines in population abundance without a compensatory decrease in disease prevalence. We apply a stochastic, spatial model of disease spread that accommodates spatial habitat heterogeneity to investigate how the population dynamics depend on multimodal (frequency-dependent and density-dependent) transmission. We found that the observed rate of population decline can be readily explained by multimodal transmission, but is unlikely to be explained by either frequency-dependent or density-dependent mechanisms alone. Multimodal disease transmission rates high enough to explain the observed decline predicted that eventual local extinction of the host species is highly likely. Our results add to a growing body of literature showing how multimodal transmission can constrain species distributions in nature.

**Open Research:** All scripts associated with the analyses in this manuscript, as well as the data we collected, are available at https://github.com/uricchio/antherSmutDis.

## Introduction

Theoretical models suggest that pathogens can limit species’ ranges, but we have relatively little empirical evidence that mechanistically connects diseases to range limits and constraints on population abundance. Many leading ideas in ecology have developed in a context where infectious diseases are considered to play a minor role in explaining species’ distribution. Pathogens and parasites are often disregarded in theories of trophic structure and ecosystem function in spite of such life-forms being more speciose and as common as their free-living counterparts (Dobson et al., 2008). Some widely-discussed exceptions exist – the potential importance of pathogens in generating species diversity, especially in tropical forests, has been frequently posited (Janzen, 1970; Connell, 1971). Large scale die-offs of in natural populations are well documented in both plants and animals and known to have transient effects on abundance (*e*.*g*., chestnut blight, Anagnostakis 1987; *Phytophthora* outbreaks, Hausbeck and Lamour 2004), but until recently many natural history observers might discount parasites and pathogens as major regulators of species distribution in favor of competition and local adaptation to environmental variables.

Despite several examples of pathogens impacting host population sizes and multiple theoretical models that posit a role for pathogens in population regulation, it is unclear whether diseases generally play a role in limiting or altering the distribution of their hosts. The “classical” view of disease would argue that pathogens, if they are host-specific, should have a negligible effect on distribution because as host abundance declines towards a margin, the population should fall below a critical threshold density for pathogen persistence. A prediction from this would be that, if range limits are accompanied by declining host densities, the edges of species ranges should be characterized by disease-free zones. And as such, disease should have little impact on range limits. Consistent with this prediction, there are empirical examples of disease prevalence (the fraction of diseased hosts) decreases near margins (Brewer, 1995; De Bellocq et al., 2002; Alexander et al., 2007). In contrast, other empirical examples suggest increases in disease prevalence near range limits (Briers, 2003). A recent empirical study of anther-smut disease (which we study herein) did not find support for a disease-free halo near range limits (Bruns et al., 2019).

This classical view, however, fails to account for at least four major factors. First, it is well appreciated that sexually-transmitted and vector-borne diseases may spread as a function of the frequency rather than density of infected individuals (Getz and Pickering, 1983; Lockhart et al., 1996). Diseases that are spread through both frequency-dependent and density-dependent mechanisms (which we term ‘multimodal’ transmission) could have especially severe effects on species’ abundance. Second, diseases that cause sterility have a much more severe effect on population size than diseases which cause mortality. Many sterilizing diseases have little effect on mortality, and therefore the duration of the infectious period is not reduced by premature death of the host (Anderson and May, 1981). Long-lived, diseased individuals can serve as reservoirs that maintain the disease and infect newly established individuals. Even with classical mass-action transmission, spatially explicit models have shown that diseases with moderate or no effect on mortality can drive host-population extinction (Boots and Sasaki, 2002; OKeefe and Antonovics, 2002), whereas this is difficult in diseases that kill their hosts. Third, when disease leads to large reductions in population size, demographic stochasticity may lead to extinctions (De Castro and Bolker, 2005) and decreased site occupancy in metapopulations (Antonovics et al., 1997). The situation may be aggravated in disease systems, where non-linear interactions lead to stochastically induced resonant cycles (Bartlett, 1957; McKane and Newman, 2005; Alonso et al., 2006) or limit cycles (Anderson and May, 1981). Fourth, habitat quality and other a/biotic factors that affect species’ range limits are likely to vary substantially over species ranges, with lower quality expected near limits (Hargreaves et al., 2014). Prior work explored the first two factors (multimodal transmission and sterility) in a mean-field model of anther-smut disease (Bruns et al., 2017), finding that the disease was unlikely to result in local extinction of populations. Spatial models that include variation in habitat quality could result in a different outcome due to the compounding effects of declining abundance (increased stochasticity) at range limits and frequency-dependent transmission that does not decline with abundance.

Here we investigate the occurrence of a sterilizing disease (anther-smut caused by the fungus *Microbotryum violaceum*) across spatial variation in the abundance of its host plant *Dianthus pavonius* over a thirteen year period. We specifically examine how the local pattern of disease prevalence and host abundance jointly depend on the spatial structure of habitat quality and transmission mode of the disease. We hypothesized that spatial variation in habitat quality combined with local reproduction might result in more severe effects of disease transmission on population abundance than in the mean-field case. Previous studies in this area have shown that the disease can reach very high levels of prevalence (*>* 40%), and that both vector and aerial transmission are important in disease dynamics (Bruns et al., 2017, 2019). We use spatially explicit models based on parameters derived from an eight-year study of marked plants (Bruns et al., 2017) to explore the factors determining local spatial patterns of disease and host distribution. Our results suggest that disease follows rather than determines relative host abundance spatially (*i*.*e*., the spatial pattern of relative abundance is determined in large part by habitat quality), but constrains total abundance across the patch. Our analysis also suggests that both aerial and vector-borne transmission rates may exceed the mean-field estimates of a previous study. Simulations of disease epidemics with transmission rates high enough to explain the observed population decline often resulted in population extinction, suggesting that anther-smut disease transmission may constrain both the abundance and distribution of *Dianthus pavonius*. We discuss a range of limitations and extensions of our analyses, including the potential for direct statistical inference of transmission parameters from the spatial disease patterns and potential deviations from our model predictions due to the effects of vector behavior and perception.

## Materials & Methods

### The study system: Anther-smut disease

Our studies were carried out on anther-smut disease caused by the fungus *Microbotryum violaceum* s. l. (*Microbotryales, Basidiomycota*) infecting the alpine carnation *Dianthus pavonius* Tausch (= *D. neglectus* Lois). *Dianthus pavonius* is a slow-growing herbaceous perennial that is endemic to the French and Italian western alps. It is found from 1500m to 2400m in grazed subalpine and upper alpine regions, flowering from July to mid-August. Anther-smut disease occurs throughout the elevational range of *D. pavonius*, with a higher incidence (% populations infected) and prevalence (% disease within populations) at lower elevations (Bruns et al., 2019).

At the vegetative stage, infections by the fungus have small but barely detectable effects (Antonovics et al., 2018), whereas at the flowering stage it sterilizes the plants by replacing the pollen in the anthers with fungal spores and preventing ovary development. Such symptoms are only expressed after the fungus enters the plant and colonizes newly developing floral buds (Schäfer et al., 2010). The disease has a long latent period. Specifically, in *Dianthus* diseased flowers are not seen until the season following infection. Infection is systemic and in most host species, including *D. pavonius*, recovery is absent or very rare (Bruns et al., 2017). The disease can be spread through at least two transmission routes: pollinator-mediated (frequency-dependent) and aerial (density-dependent) spore dispersal.

### Study site

This research was carried out near Rifugio Garelli, an alpine hostel at 2000m elevation in the Parco Marguareis (formerly Parco Naturale Alta Valle Pesio, and now administratively part of the Parco Alpi Marittime) in the Piemonte province in north-western Italy, on the northern slope of the Ligurian Alps. The general ecology, historical land-use and geology of this area is described in Gallino and Pallavicini (2000). Species names follow this publication, and Laubert and Wagner (2000).

Our study is concerned with the specific effects of a pathogen on abundance, so we were naturally concerned with other factors that might constrain population growth, such as other plants that might harbor the pathogen or effects of grazing. While there were occasional plants of species that have been found diseased with *Microbotryum* elsewhere (*Silene nutans, Silene vulgaris*, and *Lychnis flos-jovis*), none were observed to be diseased within at least several hundred meters of the study site. Farmers in the valley had grazing rights, and the area was grazed by cattle usually in late August or early September. There was no evidence of grazing of flowering plants at the time of the census. However, grazing on maturing fruits could not be excluded, and the fruits of *D. pavonius* were attacked by caterpillars or eaten by grasshoppers.

To study the relationship between host density and disease prevalence at a local scale, detailed mapping studies were carried out along a transect between 2005 and 2018, and a schematic of the transect dimensions is shown in Fig. 1. This site was chosen because it showed large gradients in host abundance, and high disease levels typical of the area.

**Fig. 1:**
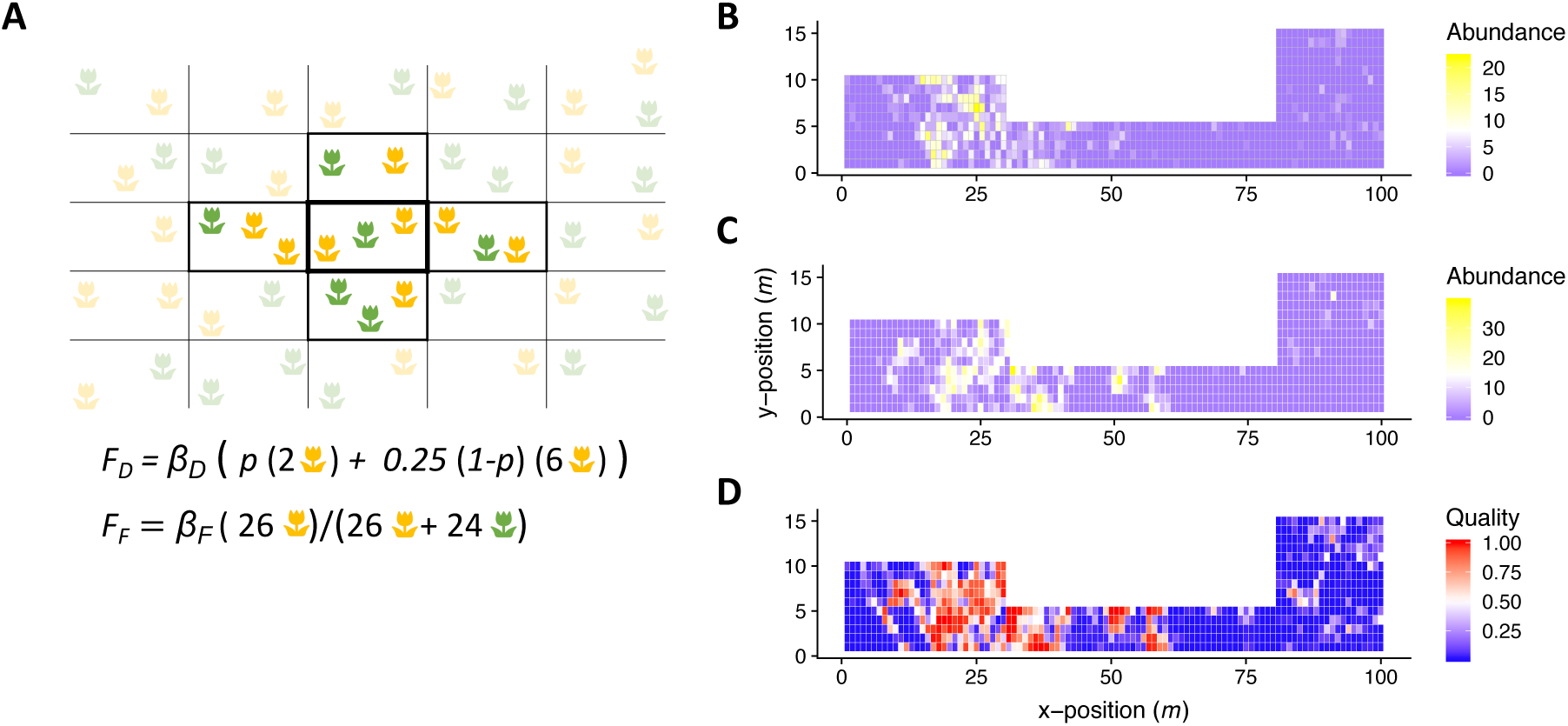
We model population dynamics on a spatial grid that captures variation in habitat quality, plant density, and disease transmission. A) Pictorial representation of the transmission model. Diseased plants are shown in yellow, healthy plants in green. Plants contributing to local density-dependent transmission are shown in darker colors, while all plants contribute to frequency-dependent transmission. *p* is the fraction of infectious spores that remain within the quadrat in which they originated, while the remaining (1 *- p*) are spread into the four adjacent transects (Fig. 1A). The force of infection experienced by the healthy plant in the central quadrat is therefore proportional to 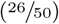, where the first term is due to the diseased plants in the same quadrat and the second is due to the six diseased plants in the neighboring quadrats. The force of infection due to vector-borne transmission is proportional to the prevalence of disease 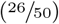. B) Observed population abundance for the transect in 2007. C) Example of population abundance in one simulation replicate. D) Empirical estimate of habitat quality for each quadrat of the transect.

The mapping area in the transect was 100m long but differed in its width in the lower (10m), middle (5m) and upper sections (15m). This widening of the transect was performed intentionally such that sample sizes would be more comparable across the length of the transect. The transect spanned an area bounded at both ends by Krummholz, a thick shrub matrix composed of *Juniperus nana* (dwarf juniper), *Rhododendron ferrugineum* (alpenrose), and *Vaccinium uliginosum* (a shrubby blueberry), in which *D. pavonius* did not occur. At the lower edge, the Krummholz was dominated by *J. nana* and *R. ferrugineum*, while the upper edge was dominated by *J. nana* and *V. uliginosum*. At the upper end, the density of *D. pavonius* decreased and there was increasing cover of *J. nana* and *V. uliginosum*; *D. pavonius* occurred as scattered individuals for another 100m above the upper edge of the transect. The open meadow between these upper and lower areas was dominated by grasses (*Festuca varia, F. pratensis, Avenella flexuosa, Agrostis schraderiana, Anthoxanthum alpinum, Phleum rhaeticum*, and scattered plants of *Festuca paniculata*). The meadow had a high diversity of forbs. At the time of flowering of *D. pavonius*, other species in flower were predominantly *Campanula scheuzeri* (blue flowers), *Potentilla grandiflora* (yellow flowers), and less commonly *Achillea millefolium* (white flowers), *Hypericum richeri* (yellow flowers) and *Alchemilla decumbens* (yellow/green flowers). The flowers of *D. pavonius* are a deep pink color, and no other species of this color were in flower, except for a small stand of *Epilobium angustifolium* (fireweed) in a patch of *Juniperus nana* where *D. pavonius* was absent.

### Transmission of anther-smut disease

Transmission can occur through at least two distinct mechanisms; pollinator-mediated or aerial. Pollinators visit diseased plants where they pick up spores and can directly deposit them on the healthy flowers, with strong analogies to vector-transmission. This mode of transmission is likely to be frequency-dependent rather than density-dependent because pollinators can adjust their flight distances in response to lower densities (Antonovics et al., 1995). Prior studies in other anther-smut systems have also confirmed that pollinator-mediated transmission is frequency-dependent (Roche et al., 1995). It is important to note that this frequency-dependent pollinator-mediated transmission route can only spread disease to flowering plants, as pollinators are unlikely to visit vegetative plants.

Aerial transmission occurs through passive shedding of the spores through either wind or water to the area close to the plant. This mode of transmission is density-dependent and local spore deposition close to source plants has been documented for several species of anther-smut (Alexander, 1990). Critically, spores that are transmitted through the aerial route may come into contact with vegetative plants, including seedlings which are known to be highly susceptible (Bruns et al., 2017).

### Population census and demography

The transect was scored for the number of diseased and healthy plants over a period of several years. Data on the presence of Krummholz and number of plants were collected for each 1m^2^ quadrat. The data were recorded in terms of both inflorescence number (each inflorescence usually had two flowers), and number of individuals. Some judgement calls had to be made when identifying separate individuals in dense patches. We excluded vegetative plants from the counts, because it is not straightforward to determine disease status in vegetative individuals. Each quadrat was also scored for percent of Krummholz coverage.

### Data analyses

We used the R package mgcv (version 1.8-27) to fit smoothed curves to the prevalence, Krummholz, and density of plants per m^2^ in each of the census years. The package allows for the fitting of generalized additive models to the observed data along the E-W transect. To perform the fitting, we used 30 knots for each year, with the exception of 2005 which only covered a subset of the transect and used 15 knots. We set the gamma parameter to 0.25. Model fitting for density, Krummholz, and prevalence used Poisson, Gaussian, and binomial errors, respectively. These data and model fits are reported in Fig. 2 (note that Krummholz coverage data was not collected in 2018).

**Fig. 2:**
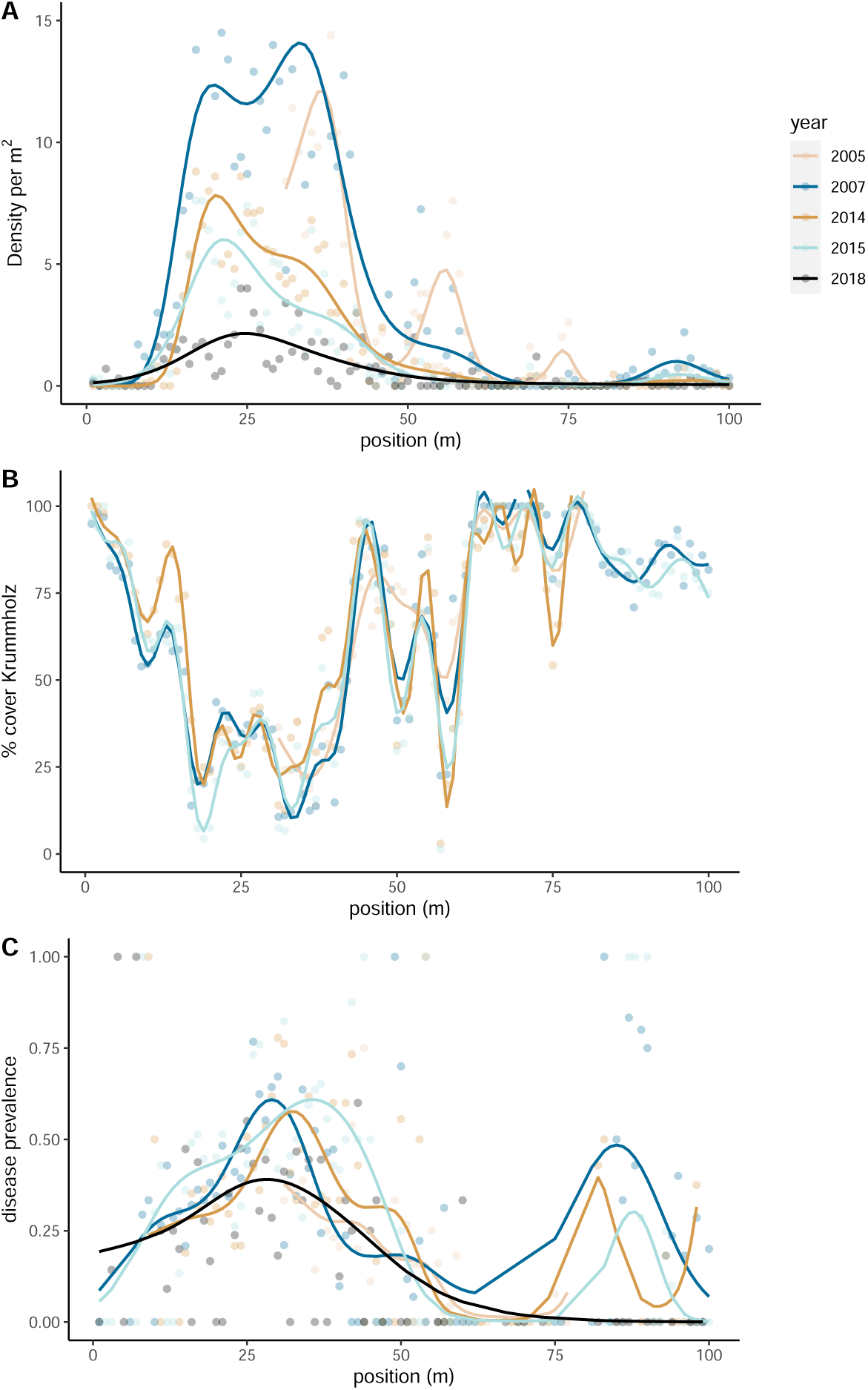
Plant density, percent cover by Krummholz, and disease prevalence (fraction of population diseased) across the transect, averaged over E-W quadrats. Model fits are described in the Methods section.

We fit quadratic (Fig. 3) and linear (Fig. 4) curves to the relationship between density and Krummholz as well as disease prevalence and density. We used the method “lm” in R for these fits. In general, when we show the data along the transect we average over the depth of the transect and show only the results along the main axis of variation (running from 0 to 100m).

**Fig. 3:**
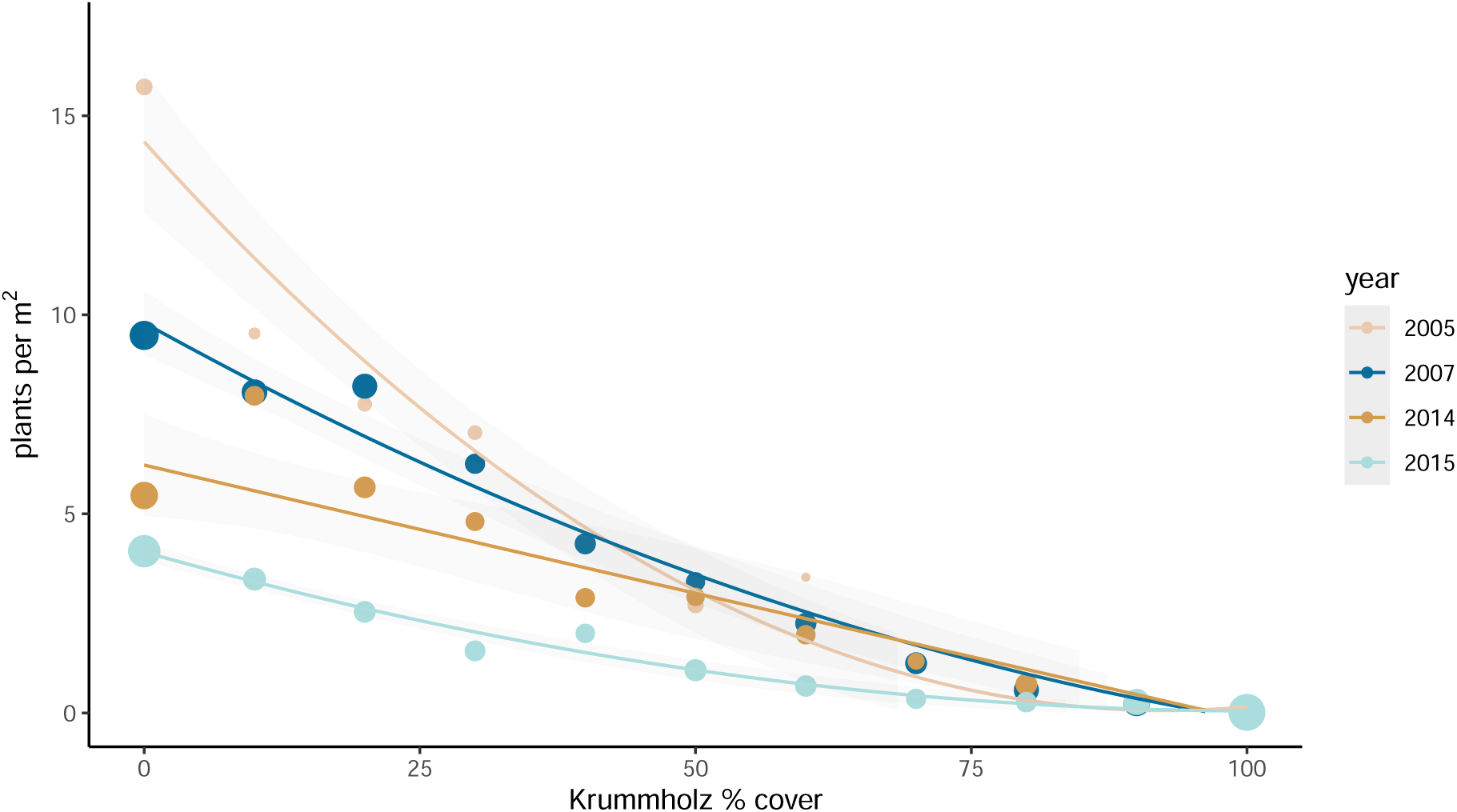
Relationship between plant density and % cover by Krummholz for different census years of the transect. Data are plotted by Krummholz class, rounded to the nearest 10% where the size of the circles is proportional to the numbers in each class. Fitted regressions are on class means with linear and quadratic components, Gaussian error, and weighted by the number in each class. The plotted fits were made using the method “`lm`” in `ggplot2` version 3.2.1. The quadratic term was significant (*P <* 0.02) in all years except for 2014. Data for 2005 are for the middle section only, and data for 2014 are for the lower and middle sections only.

**Fig. 4:**
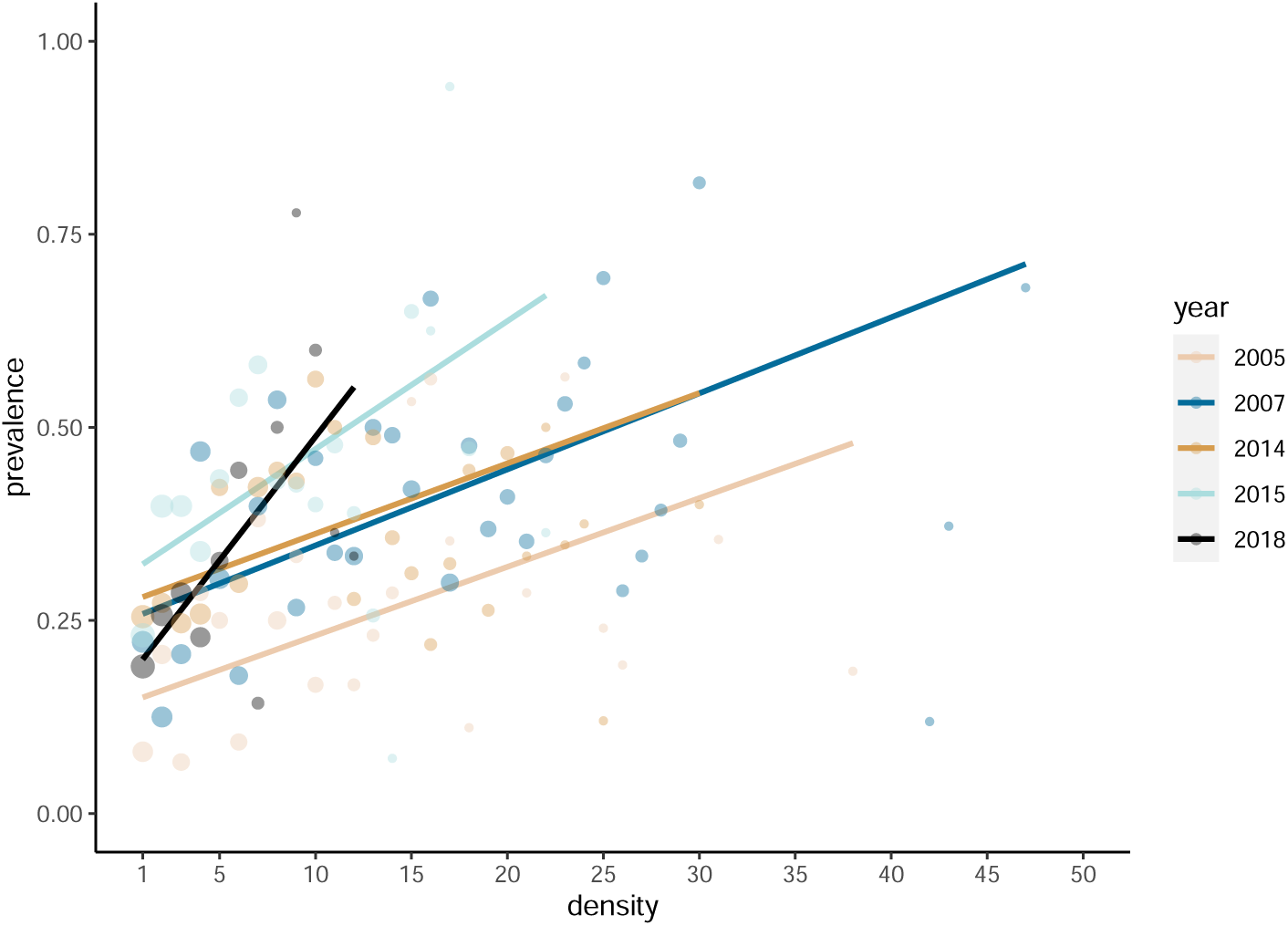
Prevalence (fraction of population diseased) as a function of density (number of plants per sq. m) for census years. Linear regression fits were made to the mean prevalence among all quadrats at each density using the method “`lm`” in `R`, weighted by the number of datapoints at each density. Circles show average values for each density class, with the size of the circle proportional to the number of quadrats at that exact density. Intercept terms and slopes were significantly different from 0 in all years (*P <* 0.02).

### Simulation methods

We extended the mean-field model of Bruns et al. 2017 to a spatially explicit grid and developed a simulation framework to generate disease patterns under the model. The goal of our simulations is to generate data that mimic the empirical transect data – in other words, numbers of diseased and healthy individuals within each quadrat on the grid as a function of time. The model includes transmission by both density-dependent (aerial) and frequency-dependent (vector-borne) mechanisms. A pictograph of the model is provided in Fig. 1A. Our simulated transect followed the same dimensions as the transect.

Our dynamic model tracks diseased and healthy individuals in a discrete-time framework where each time step corresponds to one year. We track the number of individuals in each of five distinct states – flowering-healthy individuals (*N*_*fh*_), flowering-diseased individuals (*N*_*fd*_), vegetative-healthy individuals (*N*_*vh*_), vegetative-diseased individuals (*N*_*vd*_), and juveniles (*N*_*j*_). Equations governing each of these states are given by

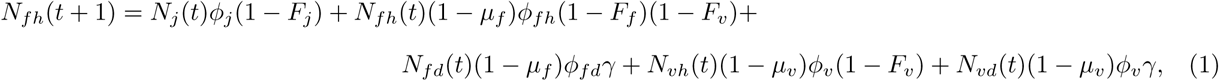

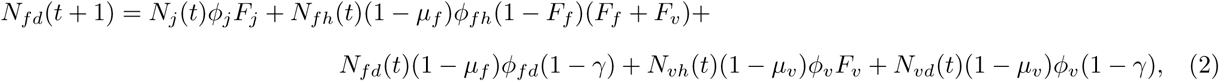

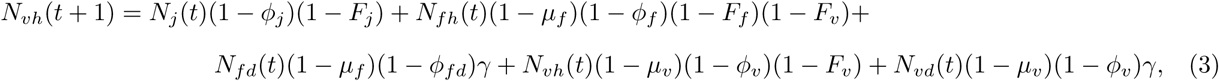

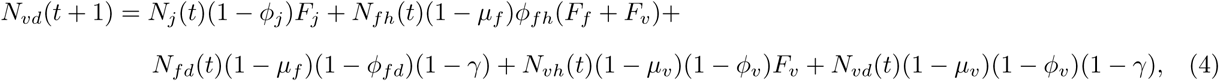

and

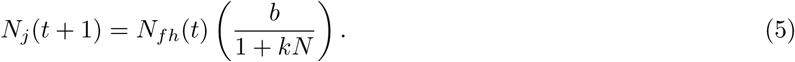

The parameters *µ, ϕ*, and *γ* represent rates of mortality, flowering, and recovery, respectively. *F* represents the force of infection. The parameter *b* gives the rate of establishment of new seedlings, while *k* is the effect of intraspecific competition on juvenile growth. Total density *N* is expressed as a sum over all five states (*N* = *N*_*fh*_ + *N*_*fd*_ + *N*_*vh*_ + *N*_*vd*_ + *N*_*j*_).

Force of infection terms control the density- or frequency-dependence of transmission. Frequency-dependent vector transmission is given by

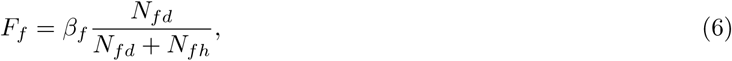

while density-dependent aerial transmission is given by

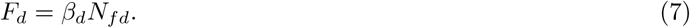

We suppose that transmission rate may differ between juveniles and adults, such that the juvenile force of infection *F*_*j*_ is given by

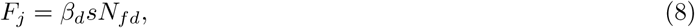

where *s* is a constant that controls the relative rate of transmission to juveniles as compared to adults. All values of *F* herein are assumed to be less than 1 such that the population sizes are constrained to be non-negative.

We adapted this model, previously developed in Bruns et al. (2017), to a spatial grid and performed stochastic simulations of transmission and reproduction on the grid (Fig. 1A). We constrain density-dependent transmission such that it is always local – individuals are infected at a rate proportional to disease frequency and/or density within the same quadrat and the four neighboring quadrats within the transect. This means that the *N*_*fd*_ terms in equations 7 and 8 are determined by the number of individuals in the local neighborhood of a focal plant, and not by the total density across the whole transect. For density-dependent transmission, we suppose that some proportion of aerial spores (*p*) stay within the quadrat where they originated, while the remaining (1 *- p*) are distributed equally over the four adjacent quadrats. Likewise, proportion *c* of seeds are retained within the origination quadrat, while (1 *- c*) are spread equally over the adjacent quadrats. Frequency-dependent transmission is assumed to be dependent on the frequency of disease across the whole transect (*i*.*e*., the term 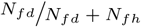 in equation 6 includes all diseased and healthy plants across the entire transect).

The number of diseased individuals within each quadrat at time *t* + 1 are sampled randomly from a Poisson distribution, in which the Poisson parameter is set equal to the expected number of individuals within each class of individuals (*e*.*g*., the expected number of individuals in class *N*_*fh*_ is given by equation 1, and the actual number is sampled from a Poisson distribution with this mean). Note that this is an approximation to the full dynamics, in which survival or transitions between diseased/healthy might be sampled binomially while reproduction might be modeled as a Poisson process – we approximate these dynamics with a single Poisson that represents the sum of each constituent process.

In addition to the dynamics governed by eqns. 1-5, our field surveys clearly demonstrated that Krummholz also plays a major role in driving the abundance of *D. pavonius*. We therefore allow habitat quality to vary proportionally to Krummholz occupancy in our simulations. We suppose that only birth rate *N*_*j*_(*t* + 1) is reduced proportional to habitat quality. A pictorial representation of the model is given in Fig. 1A, and Python code implementing our simulation method is available at https://github.com/uricchio/antherSmutDis.

### Patterns of disease prevalence and abundance

In order to establish baseline theoretical expectations, we first simulated an idealized model in which habitat quality varies from 0 to 1 over the linear transect as a simple function of position. We assumed that the transect consists of a 2-D array of quadrats with *x*-positions from 0-100. We set habitat quality (which is equivalent to percent cover by Krummholz in our field observations) to 0 for *x ∈* [1, 10] and *x ∈* [90, 100]. Transect quality increases linearly to 1 at *x* = 50, and then decreases linearly until *x* = 90. Mathematically, we multiply the patch quality by the parameter *b* (which controls the rate of production of juveniles from flowering adults). In the absence of disease, the habitat quality is effectively a linear correction on the carrying capacity of the patch.

We use the parameters inferred in Bruns et al. (2017) to run these simulations (values reported in Table 1). We ran 100 replicate simulations for each parameter combination. We stopped the simulations 50 generations after the introduction of an epidemic – this value is arbitrary, but these simulations are intended only to compare qualitative features of the models. When only frequency-dependent or density-dependent transmission were modeled, we increased the underlying transmission rates from the mean-field model such that the overall disease prevalence was similar across all three models (*i*.*e*, frequency-dependent, density-dependent, and multimodal all had final prevalence of ≈20% at 50 simulated years – to achieve this we increased the frequency-dependent transmission rate by a factor of 6 and the density-dependent rate by a factor of 2.5 relative to the values reported in Tab. 1). When exploring a range of density-dependent transmission parameters, we change *β*_*d*_ and *sβ*_*d*_ (i.e., the rate of density-dependent transmission for juvenile plants) simultaneously by the same factor (relative to the values reported in Tab. 1).

**Table 1:** Parameter values used in our study, as inferred by Bruns et al. 2017. Parameters that correspond to density-dependent processes have been scaled to accommodate the difference in spatial sacle between the studies – our study uses 1m^2^ plots, where as Bruns et al. 2017 used a 250m^2^ area. The values of *p* and *c* were not previously inferred, but our simulations suggest that population dynamics do not depend heavily on these parameters.

| Parameter definitions and values |  |  |
| --- | --- | --- |
| Symbol | Definition | Value |
| $\beta_f$ | Frequency-dependent transmission rate | 0.069 |
| $\beta_d$ | Density-dependent transmission rate for adult plants | $0.0000338 \times 250$ |
| $s\beta_d$ | Density-dependent transmission rate for juvenile plants | $0.000338 \times 250$ |
| $\gamma$ | recovery rate for diseased plants | 0.038 |
| $\mu_f$ | Mortality rate for flowering plants | 0.116 |
| $\mu_v$ | Mortality rate for vegetative plants | 0.258 |
| $\phi_{F_h}$ | Flowering rate for healthy flowering plants at $t + 1$ | 0.664 |
| $\phi_{F_d}$ | Flowering rate for diseased flower plants at $t + 1$ | 0.704 |
| $\phi_v$ | Flowering rate for vegetative plants at $t + 1$ | 0.356 |
| $\phi_j$ | Flowering rate for juvenile plants at $t + 1$ | 0.17 |
| $b$ | Establishment rate | 1.3 |
| $k$ | Effect of competition on seed output | $0.00015 \times 250$ |
| $p$ | Proportion of aerial spores remaining in quadrat | 0.9 |
| $c$ | Proportion of seeds remaining in quadrat | 0.9 |

### Estimating the rate of population decline from simulations

We sought to characterize our simulations in terms of the maximum observed rate of population decline over the time course of an epidemic. Rather than simply calculating the maximum year-over-year change, we instead fitted a logistic curve to each simulated time-series of population abundance, and calculated the maximum slope of the best-fit curve. We used the function nls (non-linear least squares) in R to perform the fitting. Since we do not have data on the number of vegetative plants in the transect, we included only simulated flowering (both diseased and healthy) plants in this calculation. We compared the results of these simulations to the rate of decline we observed in the empirical transect data, where the maximum year over year decline in the number of flowering plants occurred between 2014 (1409 plants) to 2018 (504 plants), for a yearly decrease of 226.25 plants/year. The results of these calculations are included in Fig. 6, where we simulated 20,000 replicate simulations with a range of transmission parameters (*β*_*d*_ and *β*_*f*_). All other parameters were fixed to the values reported in Tab. 1.

**Fig. 5:**
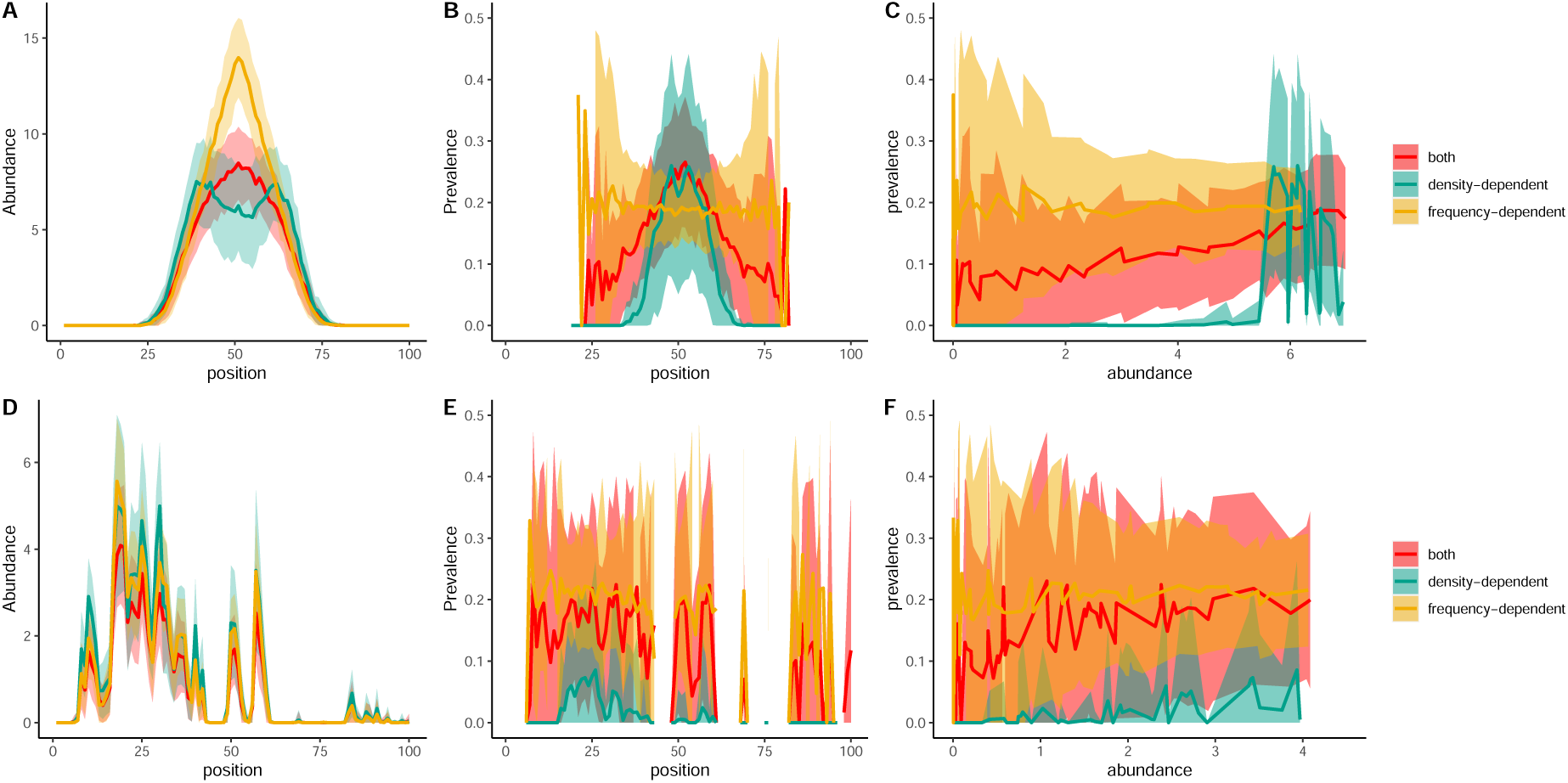
Stochastic simulations of anther-smut epidemics that used simplified spatial habitat variation, with quality declining on the edges of the transect (A-C) or realistic transect structure modeled on the observed variation in Krummholz (D-F). We report the abundance of adult plants (A&D), the prevalence of disease (B&E), and the relationship between abundance and prevalence (C&F). The dark lines represent the mean over 100 simulations, whereas the lighter colored regions represent the standard deviation of simulated outcomes (truncated at 0 on the lower edge). The multimodal transmission model (“both”) used the values of *β*_*f*_ and *β*_*d*_ reported in Tab. 1. The frequency-dependent model used a six-fold higher value of *β*_*f*_, while the density-dependent model used a 2.5 fold higher value of *β*_*d*_ such that overall disease prevalence and abundance were similar across the models.

**Fig. 6:**
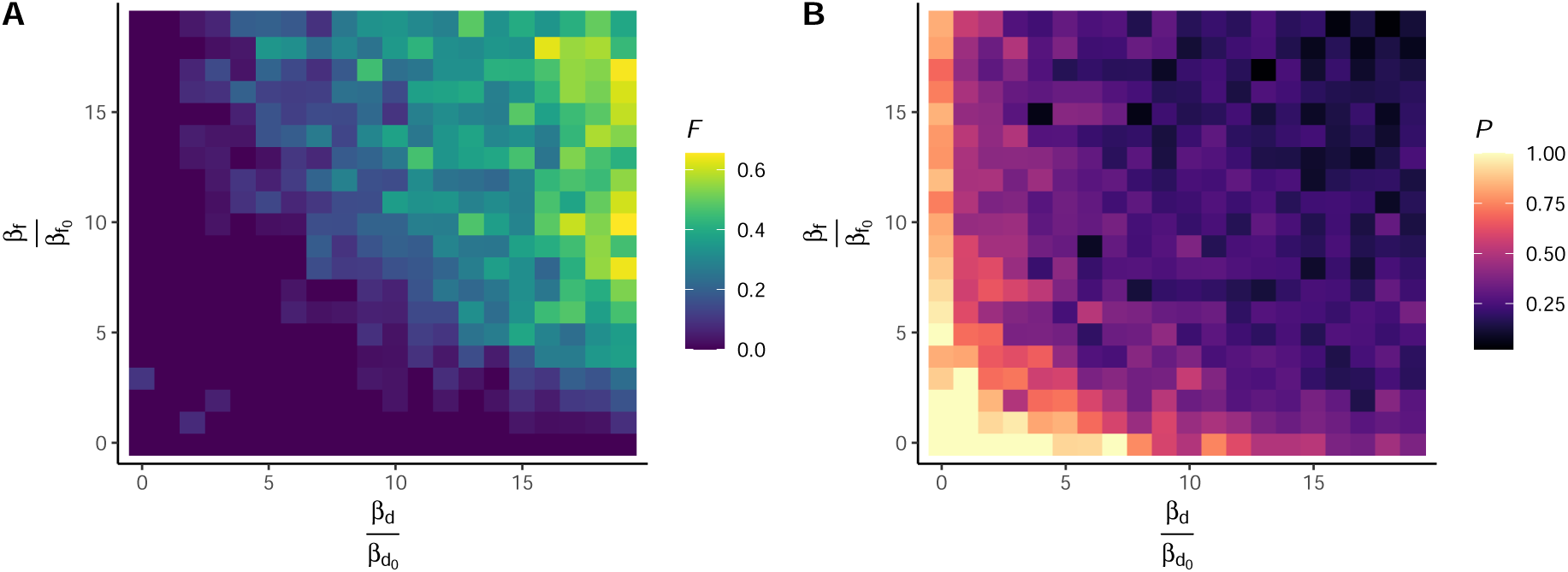
We performed simulations in which we varied the frequency-dependent (*β*_*f*_) and density-dependent (*β*_*d*_) transmission parameters relative to their mean-field estimates. In A) We report the proportion of simulations (*F*) in which the rate of decline exceeded the observed rate in the transect, and in B) we report the fraction of simulations (*P*) that resulted in local persistence (*i*.*e*., the density of plants was greater than 0 if/when the epidemic burned out of the population).

## Results

### Density and disease prevalence

There was large spatial variation in the density of *D. pavonius* (Fig. 2A) with the highest densities occurring in the 10-40m region of the transect. While this pattern was maintained over the period of the study, the number of flowering plants declined between 2007 and 2018 from over 3000 to about 500 (Fig. 2A). The spatial abundance of *D. pavonius* was strongly negatively correlated with the presence of Krummholz (*i*.*e*., dwarf juniper), not only at the transect level, but also at the quadrat by quadrat level (Fig. 3). However, the spatial distribution or percent cover by the Krummholz did not change substantially over the study period (Fig. 2B). Disease prevalence showed much less spatial variation than plant density (Fig. 2C), with prevalence being high even towards the regions of lower density at either end of the transect (with the exception being 2018, when the plants were essentially eradicated from the west end of the transect at 75-100m in Fig. 2A&C).

To further investigate the relationship between prevalence and density, we plotted prevalence against density at the quadrat level (Fig. 4). There was no evidence of a minimum density threshold for the presence of the disease – the y-intercepts and slopes were greater than zero for all census years (Fig. 4).

### Modelling spatial dynamics and transmission: theoretical expectations

Our empirical data revealed rapid population declines without substantial changes in disease prevalence (Fig. 2), but also an increasing relationship between abundance and disease prevalence in all census years (Fig. 4). We hypothesized that this relationship might be consistent with density-dependent transmission models, not frequency-dependent transmission alone. In order to clarify the expectations for the dynamics of anther-smut disease on *D. pavonius* in a spatial setting, we simulated a transect using a patch model with stochastic disease dynamics within each patch and local migration between adjacent patches (Fig. 1A). We simulated both an idealized patch quality structure in which quality increased linearly from the edges of the transect to the middle (Fig. 5A-C), and a transect modeled on the observed variation in patch quality in the transect (Fig. 5D-F). We observed that all three transmission models (frequency-dependent, density-dependent, and multimodal) resulted in abundance patterns that followed the overall patch quality, with the highest abundance near the high quality habitat in the center (Fig. 5A) or clustered near high quality peaks (Fig. 5D). However, frequency-dependent transmission alone resulted in an essentially flat relationship between position and disease prevalence, while density-dependent transmission alone resulted in a peak in prevalence near the peak in abundance (Fig. 5B). Moreover, there was a clear threshold in abundance for density-dependent transmission, with no disease present at low densities (Fig. 5C). For frequency-dependent transmission, there was no correlation between abundance and disease prevalence, while the multimodal model was intermediate. These differences between the transmission models were somewhat obscured when simulating the more complex habitat structure that mirrored the real transect, suggesting that simple density/abundance relationships will not be sufficient to distinguish transmission modes in our empirical data (Fig. 5E-F). However, density-dependent transmission alone still resulted in near-0 disease prevalence at low host plant density even in the more complex habitat structure (Fig. 5F).

### Explaining the rapid population decline

We noted that our simulations seemed to rarely (if ever) predict the high rate of population decline that we observed in the transect over the 13 year study, which was as high as a loss of 226 adult flowering plants per year. We hypothesized that increasing the transmission parameters to values higher than their mean-field estimates from Bruns et al. 2017 could result in comparable rates of population decline to those we observed. We simulated epidemics over a wide range of parameter combinations, and calculated the maximum rate of population decline for each simulation. We report the fraction of simulations that resulted in a maximum rate of population decline exceeding the observed rate (*F*, Fig. 6A). We never observed rates of decline exceeding the observed rate for simulations in which both *β*_*f*_ and *β*_*d*_ were less than 5*×* as large as their mean-field estimates (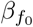 and 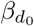). Interestingly, increasing only *β*_*f*_ or only *β*_*d*_ did not result in fast population declines, suggesting that multimodal transmission increases the potential for rapid population decline as compared to frequency-dependent or density-dependent transmission alone.

### Constraints on population abundance imposed by sterilizing pathogens

Given that the population decline cannot be explained by our simple extension of the mean-field model that uses the previous estimates of transmission, we sought to understand the implications that higher transmission rates would have for the population. When transmission was high enough to explain the observed rate of population decline, epidemics frequently resulted in local extinction of the host (Fig. 6B). Note that the criterion for extinction we used here is fairly conservative, because we consider the population to have persisted if even a single healthy (vegetative or flowering) individual remains at once the epidemic dies out – some of these persisting simulated populations might die out from subsequent stochastic extinction.

## Discussion

There have been relatively few studies on how host-specific pathogens might determine host abundance across variation in habitat quality. Parasitoids that disperse further than their hosts can restrict hosts to patches of high productivity (Hochberg and Ives, 1999). It has also been argued that increased host productivity decreases regions of stable co-existence in predator-prey systems (Rosenzweig, 1971). Beddington 1975 emphasized the potential importance of parasitoids in determining range limits. Nevertheless, the possibility that pathogens and parasites are critical determinants of range limits of host species has been the subject of limited empirical study. Here, we considered the impact of multimodal disease transmission on the spatial pattern of epidemics in alpine carnations affected by anther-smut disease.

In our empirical study, we observed that plant abundance decreased rapidly from 2005-2018, but the disease prevalence changed only modestly (Fig. 2C), a pattern that is qualitatively consistent with the predictions of frequency-dependent transmission models. In contrast, we observed that prevalence increased with density within all five census years, a pattern that is consistent with density-dependent transmission (Fig. 3). Indeed, previous work in this system has suggested that both density-dependent and frequency-dependent mechanisms are likely to operate and drive substantial disease transmission (Bruns et al., 2017). These observations led us to consider the impact of multimodal transmission on the spatial pattern of disease in anther-smut epidemics. The results of our simulations indicate that even in diseases that are completely host-specific, multimodal transmission can result in limits to host distribution, as well as signature patterns in the relationship between disease prevalence and abundance that could be help disentangle transmission mechanisms. In particular, disease prevalence is correlated with abundance as a result of density-dependence, but disease prevalence does not drop to zero at low density, likely due to frequency-dependence. We observed these patterns in both the empirical data (Fig. 4) and simulations (Fig. 5). The rapid rate of population decline that we observed is suggestive of multimodal transmission that limits host abundance and may lead to local extinction of the host. Indeed, host plants were eradicated from one end of the transect in 2018 (Fig. 2C).

Many documented examples of disease-driven extinction are characterized by reservoir species that spill-over in to more susceptible hosts (De Castro and Bolker, 2005; Mordecai, 2011). In contrast, studies on diseases that are host-specific are less prevalent, perhaps because of the presumption that disease should fade out at low density where transmission should be lower. Even highly lethal pathogens are predicted to be ineffective drivers of extinction, because the fact that they kill their hosts decreases the duration of infection and therefore leads to a diminishing impact on host population size (Anderson and May, 1981). Diseases that cause sterility are widespread in plant pathogens (Clay, 1991; Wennström et al., 2003) and in animal parasites (parasitic castration; Kuris 1974; Baudoin 1975; Jaenike 1992), and sterilization also characterizes many sexually transmitted diseases (Lockhart et al., 1996). Our results suggest that multimodal transmission of a sterilizing disease, which could be attributed to simultaneous vector-borne and aerial spore deposition, can result in very fast population declines and limitation of host abundance and range.

Transmission rates of frequency-dependent and density-dependent transmission that were high enough to explain the high rate of decline that we observed often resulted in population extinction in our simulations (Fig. 6). In contrast, frequency-dependence or density-dependence alone rarely resulted in rates of decline as high as those we observed, even with rates of transmission up to 20 times as high as were previously inferred. This may be due to the complementary effects of frequency-dependent and density-dependent transmission. Density-dependence alone results in slow transmission in low density patches, in which plants experience a very low force of infection. Frequency-dependence alone results in essentially uniform rates of transmission across the transect, meaning that high-density areas decline no faster than low density areas. In a well-mixed population model without habitat quality variation, this distinction is not possible, because the rate of local transmission and global transmission are the same. When frequency-dependence and density-dependence are combined, infections in our model had greater likelihood of being sustained and causing local extinction.

It should be noted that there are some limitations of our simulation framework and modeling approach. First, the model that we simulate is parameter-rich and exploring this parameter space fully would be computationally prohibitive. While the experimentally-determined demographic parameters in our simulations can generate spatial epidemic data that closely mimic the observed data in the transect, it is possible that changes in some of these parameters (for example, those pertaining to competition and establishment) could affect the disease dynamics and change predicted population outcomes. We were particularly concerned that seed and spore dispersal rates higher than those we considered could make substantial differences in the outcomes, especially if low-density patches could experience higher force of infection through proximity to higher-density patches. Although increasing the rate of spore and seed dispersal did slightly alter the relationship between the rate of population decline and disease outcomes, it did not qualitatively affect any conclusions of our study (Fig. S1). Second, while the model that we proposed captures the demographic rates for important life history components in *Dianthus*, it makes several simplifying assumptions about transmission. In particular, density-dependent transmission is assumed to occur locally only in adjacent quadrats, meaning that aerial transmission is limited to an ≈2m range, as supported by previous empirical work (Bruns et al., 2017). Frequency-dependent transmission occurs completely independent of local prevalence and abundance in our model, implying that pollinators move between plants in a manner that is completely independent of plant density. In practice, both of these assumptions are likely to be violated to some degree, with some longer-range aerial transmission and some dependence of vector movement and decision-making on local plant density. Third, our modeling does not account for other processes that may act to limit the range and abundance of alpine carnations, such as herbivory and environmental change.

Despite these limitations, our study captures the extremes of both density-dependence and frequency-dependence in disease transmission, and shows how multimodal transmission can contribute to elevated rates of population decline in anther-smut epidemics. Our work suggests that greater consideration should be given to transmission dynamics of sterilizing diseases and the impact they may have in determining host abundance and species distributions in nature.

## Acknowledgements

We thank the support of the staff of the Parco Naturale del Marguareis, especially Valentina Carasso, Ivan Pace and Bruno Gallino. We also thank Guido and Adriana Colombo, for their hospitality at the Rifugio Garelli. Additional field assistance was provided by Colin Antonovics, Ben Adams, Amy Blair, Lidia Castagnoli, Dylan Childs, Jae-Hoon Cho, Claudio Ferracciolo, Mary Gibby, Mandy Gibson, Ruth Hamilton, Ed Jones, Marika Mandaglio, Chiara Mattalia, Mike Paetz, Tim Park, and Ian Sorrell. Students assisting with the long-term census included Indigo Ballister, Audrey Batzel, Rachel Cohen, Amy Johnson, Noah Lerner, Ian Miller, Anthony Ortiz, Laura Pierce, Robbie Richards, Lisa Rosenthal, Molly Scott, Casey Silver, Adrianna Turner, Monroe Wolfe, and Sarah Yee. The early data were the result of a travel grant from the University of Sheffield to Mike Boots and Alex Best, and a University of Virginia summer assistantship to Jessie Abbate. Subsequent support to J. Antonovics, E. Bruns, M. Hood, and M. Boots came mainly from the joint NSF-NIH-USDA Ecology and Evolution of Infectious Diseases program, including DEB-1115899, DEB-1115765, R15GM119092, and R01GM122061. Student support was funded in part by the NSF REU program, and by their respective universities.

## Supplemental Figures

**Fig. S1:**
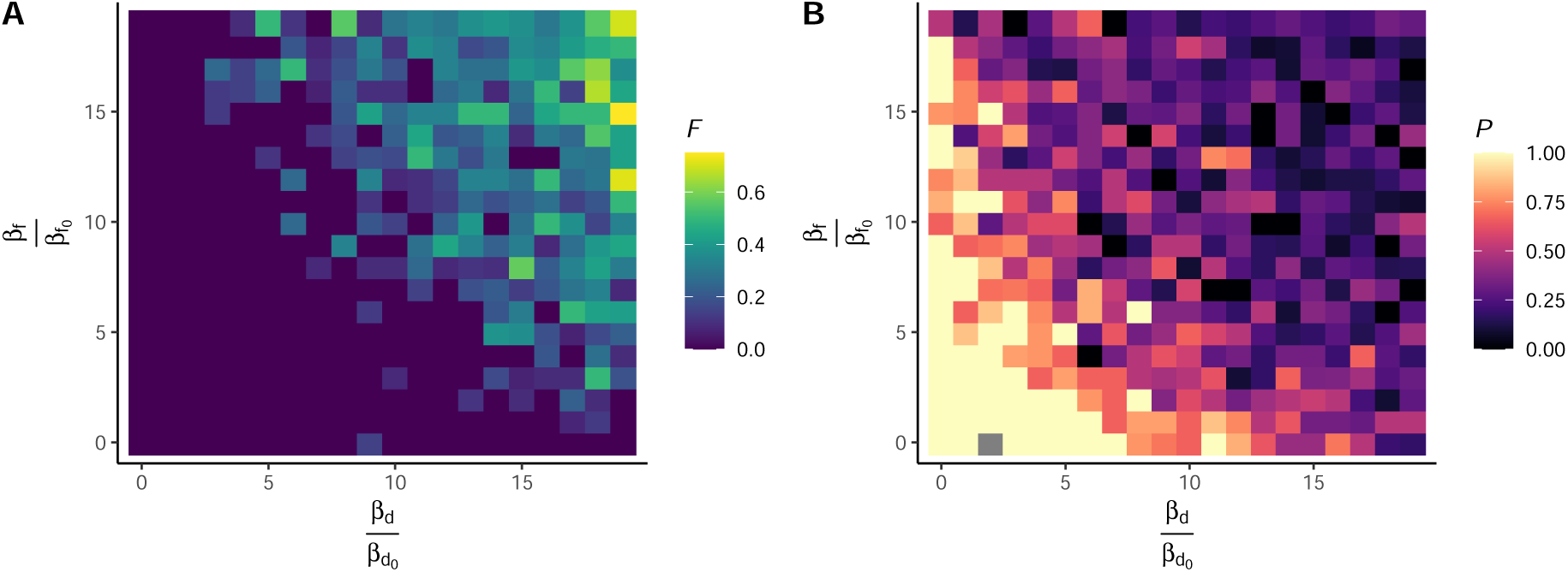
Simulations in which we varied the frequency-dependent (*β*_*f*_) and density-dependent (*β*_*d*_) transmission parameters relative to their mean-field estimates. Here we additionally set *p* = 0.5 and *c* = 0.5, meaning that only half of spores and seeds remain within the local quadrat in which they originated, and the other half are distributed equally over the other adjacent transects. In A) We report the proportion of simulations (*F*) in which the rate of decline exceeded the observed rate in the transect, and in B) we report the fraction of simulations (*P*) that resulted in local persistence (*i*.*e*., the density of plants was greater than 0 if/when the epidemic burned out of the population). Note that only 5,000 simulations were performed to study this condition, relative to 20,000 reported in Figure 6.

